# Test-retest reliability and convergent validity of (R)-[^11^C]PK11195 outcome measures without arterial input function

**DOI:** 10.1101/298992

**Authors:** Pontus Plavén-Sigray, Granville James Matheson, Zsolt Cselényi, Aurelija Jučaite, Lars Farde, Simon Cervenka

## Abstract

**Background:** The positron emission tomography radioligand (R)-[^11^C]PK11195 can be used to quantify the expression of translocator protein (TSPO), which is considered a marker for activation of glial cells. TSPO is expressed throughout the brain, and for this reason no true reference region exists. When a radioligand does not have a reference region, an arterial input function (AIF) is usually required in order to quantify binding. However, obtaining an AIF can be difficult as well as uncomfortable for participants. Alternative methods have therefore been proposed with the aim of estimating (R)-[^11^C]PK11195 binding without arterial measurements, such as standardized uptake values (SUVs), supervised-cluster analysis (SVCA), or the use of a pseudo-reference region. The objective of this study was to evaluate the test-retest reliability and convergent validity of these techniques.

**Methods:** Data from a previously published (R)-[^11^C]PK11195 test-retest study in six healthy male subjects were reanalysed. Non-displaceable binding potential (BP_ND_) was calculated for a set of cortical and subcortical brain regions using the simplified reference tissue model, with either cerebellum as reference region or a reference input derived using SVCA. SUVs were estimated for the time interval of 40-60 minutes. For comparison, total distribution volume (VT), specific distribution volume (V_S_) and BP_ND_ were estimated from the two-tissue-compartment model (2TCM) using AIF. Test-retest reliability was then assessed for all outcome measures. Convergent validity was examined by correlating all measures derived without an AIF to those derived using 2TCM.

**Results:** Test-retest reliability for BP_ND_ estimates were poor (80% of all regional ICCs<0.5). SUVs showed, on average, moderate reliability. BP_ND_ estimates derived without an AIF were not correlated with V_T_, V_S_ or BP_ND_ from the 2TCM (all R^2^<12%). SUVs were not correlated with any other outcome (all R^2^<9%).

**Discussion:** BP_ND_ estimated using cerebellum or SVCA as reference input showed poor reliability and little to no convergent validity with outcomes derived using an AIF. SUVs showed moderate reliability but no convergent validity with any other outcome. Caution is warranted for interpreting patient-control comparisons employing (R)-[^11^C]PK11195 outcome measures obtained without an AIF.

## Introduction

(R)-[11C]PK11195 was the first positron emission tomography (PET) radioligand developed for quantification of the translocator protein (TSPO). Within the brain, TSPO is mainly expressed in glial cells. Based on in vitro studies showing increases in TSPO expression in response to pro-inflammatory stimuli, the protein has been considered a biomarker for brain immune activation (Venneti, Lopresti, and Wiley 2013). As such, (R)-[11C]PK11195 has, since the early 1990s, been applied in a wide range of clinical studies (Politis, Su, and Piccini 2012).

TSPO is expressed throughout the brain which means that no part can serve as reference region in quantification of specific (R)-[11C]PK11195 binding. Instead, a metabolite-corrected arterial input function (AIF) must be obtained and used as an input function for a kinetic model from which binding estimates can be estimated. Common measures of regional binding derived from the use of an AIF are total distribution volume (V_T_), specific distribution volume (V_S_ or BP_P_) and binding potential (BP_ND_) (R. B. Innis et al. 2007).

Obtaining a metabolite-corrected input function is costly, often uncomfortable for research participants, and can also be prone to measurement error. Therefore, alternative quantitative approaches for quantifying binding have been suggested which are less demanding and which do not require an AIF. The most simple method is to calculate the radioactivity concentration in a brain region normalized by the injected radioactivity and the subject’s weight (standardized uptake value, or SUV). As such, the SUVs does not directly reflect specific binding since the signal also contain non-specific binding and radioactivity from vasculature. Importantly, SUVs are also dependent on the rate and extent of radioligand delivery to the brain. This means that results may be influenced by cerebral blood flow, or peripheral changes such as differences in metabolism or blood binding. For TSPO in particular, its high concentration in peripheral tissues, which is in turn sensitive to peripheral inflammation, can greatly influence the extent of radioligand brain delivery (Imaizumi et al. 2007). Due to these reasons, SUVs might not be a suitable index of TSPO binding in brain.

An alternative way of quantifying (R)-[11C]PK11195 binding without the use of an AIF is the supervised cluster analysis (SVCA) method (F. E. Turkheimer et al. 2007; Yaqub et al. 2012). SVCA, which is performed on dynamic PET images, aims to segment voxels into classes, differentiated by their kinetic behavior. The goal is to isolate gray matter (GM) voxels assumed to contain negligible levels of specific binding. These voxels are then used to establish a time-activity-curve (TAC) serving as a reference input in a kinetic model, such as the simplified reference tissue model (SRTM) (A. A. Lammertsma and Hume 1996). The SVCA method has been used in (R)-[11C]PK11195 studies which, for example, compared binding between healthy control subjects and patients with Alzheimer’s disease (B. N. Van Berckel et al. 2008; Parbo et al. 2017), multiple sclerosis (Rissanen et al. 2014), traumatic brain injury (Folkersma et al. 2011), schizophrenia (Van Der Doef et al. 2016), or studies which examined changes in TSPO expression in normal aging (Schuitemaker et al. 2012; Kumar et al. 2012).

Another simplified approach to obtain BP_ND_ values without arterial sampling is to use a reference tissue model with cerebellum as reference region, despite the fact that the cerebellum contains non-negligible levels of TSPO (Doble et al. 1987). This method has been used, for example, to compare (R)-[11C]PK11195 binding in healthy controls to patients with psychosis or schizophrenia (S E Holmes et al. 2016; Di Biase et al. 2017), major depressive disorder (Sophie E Holmes et al. 2018) and glioma (Z. Su et al. 2013). Since there is specific binding of (R)-[11C]PK11195 in the reference region, ensuing BP_ND_ values will not reflect the “true” binding, but rather relative regional binding to target.

In order for PET quantification methods to be useful in clinical studies they should yield outcomes which are both reliable and valid. In a previous test-retest study of six healthy subjects performed at our center, the reliability of (R)-[11C]PK11195 BP_ND_ values obtained using AIF were found to be very poor in most target regions examined (Jučaite et al. 2012). In contrast, the test-retest reliability of (R)-[11C]PK11195 BP_ND_ from SRTM with SVCA reference has been evaluated in four patients with Alzheimer’s disease (F. E. Turkheimer et al. 2007). In that study, ICC values were found to be high in most regions of interest. However, no study has yet examined the test-retest reliability of SVCA in healthy controls. To our knowledge, the reliability and convergent validity of (R)-[11C]PK11195 SUV or BP_ND_ from SRTM with cerebellum as reference have never been reported, despite both outcomes being applied in clinical studies.

The main objective of this study was to evaluate the test-retest reliability and repeatability of (R)-[11C]PK11195 1) SUVs and 2) BP_ND_ obtained from SRTM, using cerebellum or SVCA derived voxels as reference, respectively. The second objective was to examine the convergent validity of these outcomes by correlating them to V_T_, V_S_ and BP_ND_ values derived using an AIF.

## Methods and Materials

### Subjects and imaging procedures

In the present analysis we included PET examinations from six healthy male subjects (mean age = 25.8, *±* 3.9) who participated in a previous test-retest study of (R)-[11C]PK11195 (Jučaite et al. 2012). All subjects gave written informed consent according to the Helsinki declaration prior to their participation in the original study. The study was approved by the Karolinska University Hospital Radiation Safety Committee and the Regional Ethics Committee in Stockholm.

All subjects participated in two PET measurements that took place approximately 6 weeks apart, and were run on an ECAT Exact HR 47 system (Siemens/ CTI, Knoxville, TN, USA). Structural Magnetic Resonance Imaging (MRI) examinations were performed on a Siemens 1.5 T Magnetom, resulting in a T1-weighted image for each subject. Production and radio-synthesis of (R)-[11C]PK11195 has been described previously (Jučaite et al. 2012). Mean injected radioactivity was 302 *±* 33 MBq. Arterial samples were obtained in all PET measurements, from which a metabolite-corrected AIF was derived (Jučaite et al. 2012).

ROI delineation was performed on the subjects’ T1-weighted images using the FreeSurfer software (5.0.0, http://surfer.nmr.mgh.harvard.edu/). ROIs were co-registered to PET images using SPM5 (Wellcome Department of Cognitive Neurology, UK). Sixty-three minute TACs were extracted for the whole of greymatter (GM), frontal cortex, striatum, thalamus, hippocampus and cerebellum (CER), except for one PET examination were only a 50 minute scan was obtained.

### Quantification of outcomes with and without AIF

The two-tissue compartment model (2TCM) with AIF was used to estimate kinetic rate constants. The fraction of blood volume in target tissue (vB) and the delay between start of the AIF and the ROI TAC were fitted using the whole greymatter TAC. Ensuing values were held constant for the remaining ROI fits. V_T_, V_S_ and BP_ND_ were then calculated using the rate constants.

SUVs were calculated from the average radioactivity concentration in frames spanning from 40-60 minutes of the regional TACs, and dividing by the injected radioactivity and the subject’s body weight. A time span of 40-60 minutes was chosen since this has previously shown to produce SUVs which were associated with V_T_ in knee joints in patients with rheumatoid arthritis (Van Der Laken et al. 2008).

The original SVCA method classifies PET voxels into six different tissue-types associated with distinct kinetic profiles: 1. GM with high specific binding 2. GM with low specific binding, 3. whitematter, 4. soft tissue, 5. bone and 6. blood. It has been shown that removal of bone and soft tissue, by using a MRI defined brain-mask, prior to performing SVCA reduced variability of binding estimates and improved correlation to outcomes derived using an AIF (Boellaard et al. 2008). We therefore applied this restricted SVCA method (SVCA4), using the Matlab software “Super-PK” (Imperial Innovations, Imperial College London) and two different sets of population-based kinetic classes (F. E. Turkheimer et al. 2007; Yaqub et al. 2012). The Super-PK software was modified in order to be compatible with the scanning protocol applied in this study. Specifically, a cubic Gaussian smoothing kernel (FWHM 4mm) were applied to all PET images prior to the analysis, and the 30 second background frame present in the population based kinetic classes from F. E. Turkheimer et al. (2007) was removed. A reference TAC was then obtained for each PET measurement consisting of GM voxels classified as being associated with low specific binding. SRTM (called SRTM-SVCA4 below) was applied to estimate BP_ND_ for all ROIs. We also estimated V_T_ of the SVCA reference TACs using the 2TCM in order to ascertain that the results were similar to previously published data on young healthy controls. In this study we present only outcomes using the population based kinetic classes from F. E. Turkheimer et al. (2007) as these produced the most robust results.

It has also been shown that by using a version of SRTM that takes the radioactivity contribution from the vasculature into account, separation in (R)-[11C]PK11195 BP_ND_ between patients with AD and healthy controls can be improved (Tomasi et al. 2008; Yaqub et al. 2012). In addition to the SRTM algorithm, this model (called SRTMv) estimates and corrects for the fraction of blood volume in both target and reference TACs, by using an image-derived blood curve (Tomasi et al. 2008). Hence, we also evaluated the performance of SRTMv when using a reference curve derived from SVCA4 (SRTMv-SVCA4). Image-derived blood curves were obtained by extracting radioactivity from the entire scan from a region defined by the 10 voxels of highest-intensity from the first minute of each examination, as described previously (Tomasi et al. 2008).

Finally, the SRTM with cerebellum as pseudo-reference region (SRTM-CER) was also applied on all PET measurements and TACs to obtain BP_ND_ values for each ROI.

### Statistical analyses

The test-retest reliability, repeatability and precision were examined by calculation of the the intra-class correlation coefficient (ICC), the percentage average absolute variability (AbsVar) and the standard error of measurement (SEM) respectively. Since AbsVar can scale with the additive magnitude of the outcome, this particular metric is not suitable for comparing different outcomes with different means. We therefore also report the test-retest metric minimum detectable difference (MD). MD is based on the precision of an outcome (SEM) and is an approximation of the size of a difference from one measurement to another measurement which would be needed to detect a “real” change (according to a 95% confidence interval; Weir (2005)). MD is reported as a percentage of the absolute mean of the outcome, in order to allow for comparison between different measures. Convergent validity was examined by correlating all outcomes without AIF to those derived using AIF.

All kinetic modelling was performed using the R-package “kinfitr” (version 0.3.0, www.github.com/mathesong/kinfitr) together with “nls.multstart” (Padfield and Matheson 2018). All statistical analyses were carried out in R (v.3.3.2 “Sincere Pumpkin Patch”).

## Results

Table 1 shows the mean, SD and test-retest metrics for all outcomes. BP_ND_ values from SRTM using SVCA4 and cerebellum as reference, and SRTMv using SVCA4 as reference were in the same range as described previously for healthy control subjects (F. E. Turkheimer et al. 2007; Yaqub et al. 2012). There was a large difference in magnitude of BP_ND_ values derived with and without the use of an AIF. Regional BP_ND_ from 2TCM were on average 7 times higher than BP_ND_ from SRTM-SVCA4 and over 700 times higher than BP_ND_ from SRTM-CER. V_T_ values of the SVCA4 reference TACs were in the same magnitude and range (mean = 0.74, sd = 0.18, range = 0.49 to 0.96) as previosly published results (Yaqub et al. 2012).

**Table 1:**
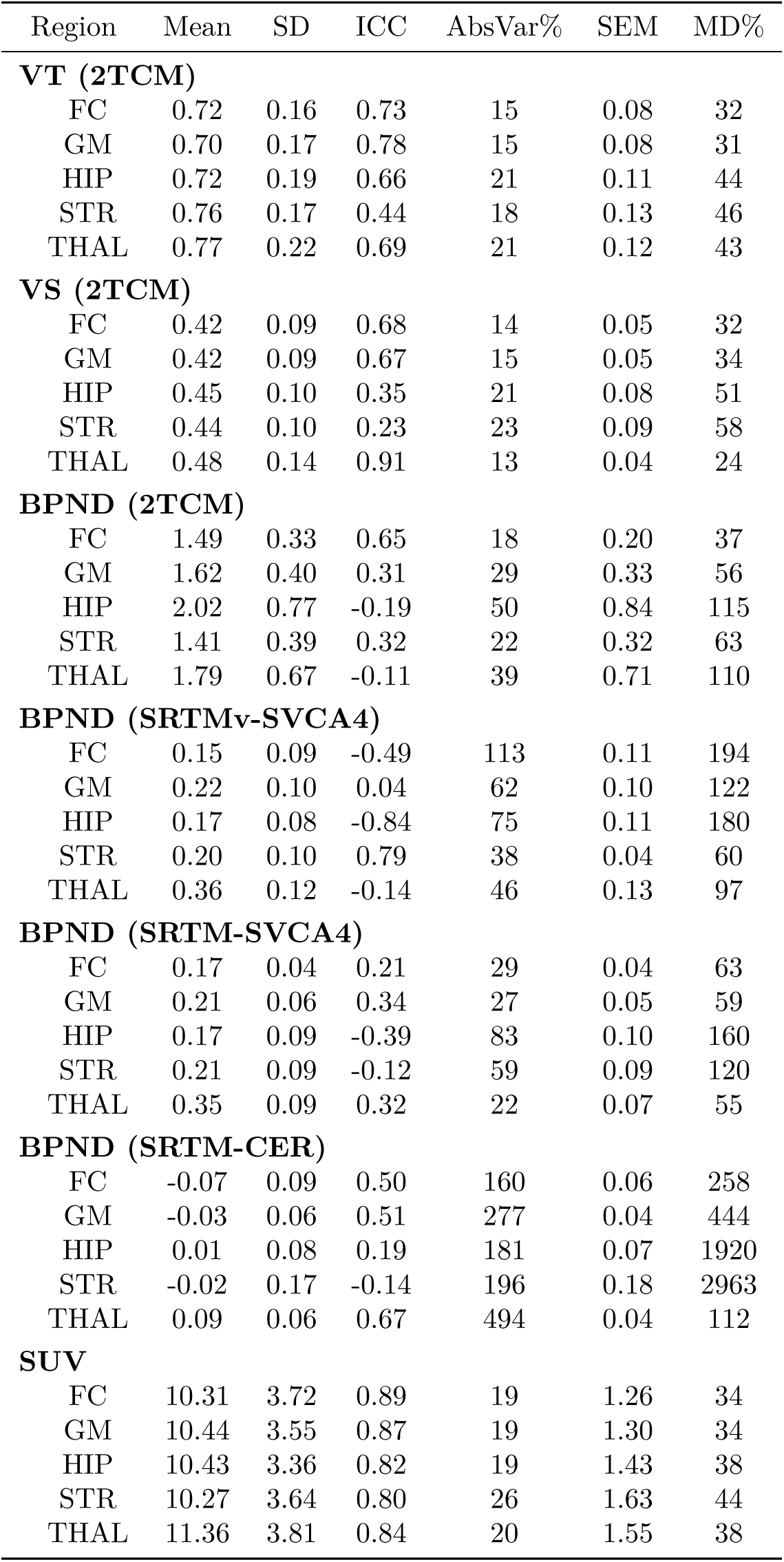
Mean values (for both PET examinations) and rest-retest reliability, repeatability and precision estimated using the Intra-Class Correlation Coefficient (ICC), average absolute variability in percentage (AbsVar) and standard error of measurement (SEM), of different outcome measures derived with or without AIF. The minimum detectable difference (MD) denotes the difference (expressed as a percentage of the mean) needed between two measurements for them to be significantly different from each other

In the present analysis, SUVs, V_T_ and V_S_ had the highest reliability across all ROIs (median ICC_SUV_ = 0.84; median ICC_VT_ = 0.69; median ICC_VS_ = 0.67). BP_ND_ from SRTM and SRTMv with SVCA4 reference showed the lowest overall reliability (median ICC = 0.21 and -0.14).

SUV, V_T_ and V_S_ showed on the lowest detectable difference (median MD_SUV_ = 38; median MD_VT_ = 43; median MD_VS_ = 34), while BP_ND_ from SRTM-CER showed the highest MD (median MD = 444).

Figure 1 shows the relationships between all (R)-[11C]PK1195 outcomes derived using AIF (V_T_, V_S_ and BP_ND_) and all outcomes derived without using AIF (BP_ND:SVCA_, BP_ND:CER_ and SUV). The correlation between BP_ND_ from 2TCM v.s. BP_ND_ from SRTM-SVCA4, SRTMv-SVCA4 or SRTM-CER was negligible to non-existent, with an explained variance < 2% for all associations. V_T_ and V_S_ were highly correlated (69% explained variance), but neither showed a strong association with BP_ND_ from AIF (both explained variances < 9%). SUVs were not correlated to any other outcome measures (explained variance < 9%).

**Figure 1:**
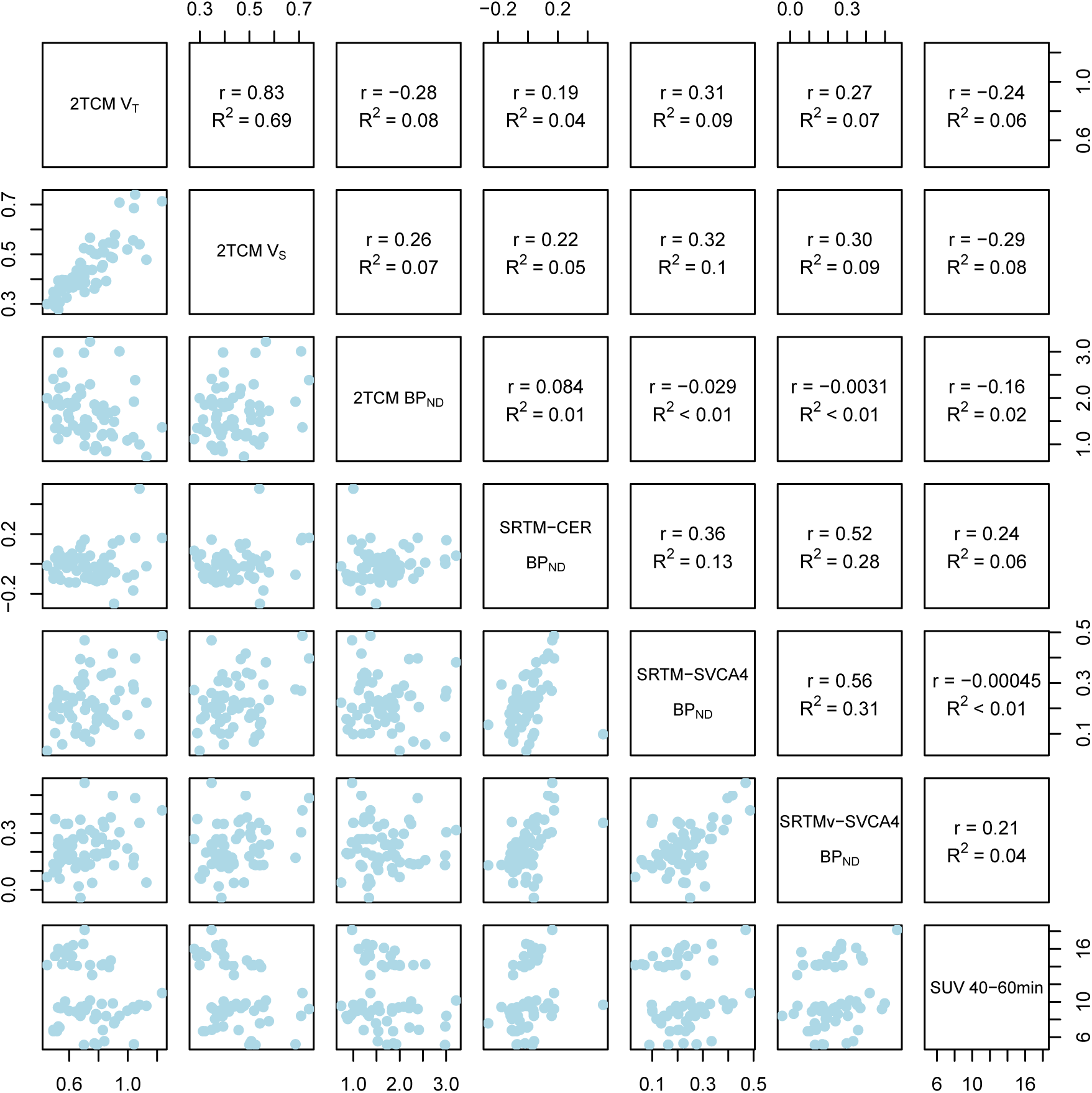
Relationships between all (R)-[11C]PK1195 outcome measures. Values from both PET examinations and all regions have been pooled in each panel. Pearson’s correlation coefficients (*r*) and explained variance (*R*^2^) are presented in the upper diagonal.

## Discussion

The objective of this study was to examine the reliability and convergent validity of (R)-[11C]PK11195 outcomes commonly applied in clinical *in vivo* studies of TSPO binding. Specifically, we evaluated outcome measures of radioligand brain exposure and binding which do not require an arterial input function (AIF), and compared them with binding outcomes derived using an AIF (i.e. V_T_, V_S_ and BP_ND_ from the 2TCM).

There was a striking difference in magnitude between BP_ND_ values from 2TCM using a AIF, and BP_ND_ values from SRTM-SVCA4 and SRTM-CER, with BP_ND_ values from 2TCM being much higher compared to other two measures. This signifies that the use of SVCA, as well as cerebellum, for derivation of a reference TAC yields only relative or pseudo-BP_ND_ values. TSPO is expressed throughout the brain, and specific binding is to be expected in every voxel (Doble et al. 1987; Farde 2015). Hence, it is unlikely that SVCA4 or cerebellum can be used to establish a TAC that reflects a true reference, devoid of TSPO, for (R)-[11C]PK11195.

In general, all (R)-[11C]PK11195 outcome measures analysed in this study showed poor to moderate reliability. For whole-greymatter only SUV and V_T_ showed, on average, acceptable reliability (ICC > 0.65), and for all regions and outcomes evaluated, only thalamus V_S_ reached the recommended threshold for clinical use (ICC > 0.90) according to previously suggested criteria (Portney and Watkins 2009). Assuming that the true TSPO concentration is stable between PET examinations, an ICC of 0.5 suggests that as much of the variance in the sample can be attributable to signal as attributed to measurement error and noise. All outcomes derived without the use of an AIF showed ICC values around or below 0.5, suggesting poor reliability for these measures. SRTM with cerebellum as reference region showed the largest imprecision and MD. This suggest that a change in BP_ND_ from SRTM_CER_ would need to be, on average, larger than 10 times the mean in order to detect a true difference between two measurements of the same subject. In comparison, a change in V_S_ of (in average) 40% would be necessary to detect a difference that is not only due to noise. One potential reason for the lack of reliability and precision for BP_ND_ from cerebellum and SVCA is that the target and pseudo-reference TACs are similar in shape and magnitude. This produces BP_ND_ values close to zero (or negative) which are sensitive to even small amounts of measurement error.

In addition to the above, the use of cerebellum as reference would also require researchers to establish significant *equivalence* (Schuirmann 1987; Lakens 2017) in reference region specific binding between the groups which are being compared. A non-significant difference between groups does not translate into evidence in favor of an absence of a difference (Dienes 2014), contrary to conclusions sometimes drawn in literature.

V_T_, V_S_ and BP_ND_ derived from 2TCM showed little to no correlation with BP_ND_ derived using outcomes without an AIF. This indicates that BP_ND_ from the reference input models have no convergent validity in relation to binding outcomes from AIF, and vice versa. Hence, if either V_T_, V_S_ or BP_ND_ derived using an AIF is to be considered an approximate index of specific TSPO binding, then BP_ND_ derived without the use of AIF cannot be considered valid. However, BP_ND_ from AIF also produced low ICC values and negligible association with V_T_ and V_S_, suggesting that this outcome is also unreliable and unstable. SUVs showed the highest average reliability but were not correlated with any other outcome measures.

In healthy control subjects, a large portion of the (R)-[11C]PK11195 signal consists of non-specific binding and unbound radioligand, as determined by blocking studies showing BP_ND_ values in the range of 0.8-0.9 (Kobayashi et al. 2017). A low signal for specific binding in healthy controls may partly explain the low reliability observed in this study. In comparison, much higher reliability has been shown for SVCA in patients with Alzheimer’s disease (F. E. Turkheimer et al. 2007) where glial cells are known to be elevated based on post-mortem studies (Heneka et al. 2015). Second generation TSPO tracers, which show higher specific binding (Fujita et al. 2017), also display higher ICC values in healthy control subjects (Collste et al. 2016). For (R)-[11C]PK11195, the low reliability means that only very large effects are possible to detect. While such effects may be present in some patient groups, such as Alzheimer’s disease, caution is advised for disorders were changes in TSPO might be more subtle.

Importantly, the 6-week interval between PET measurements in this study means that TSPO levels may change from test to retest. This, in turn, would lead to lower reliability and precision. However, since many clinical studies aim to evaluate longitudinal interventions, or correlate (R)-[11C]PK11195 outcomes with more stable independent variables, this interval mimics that of realistic and relevant designs of PET studies. In addition, the time between measurements also should not impact the relative reliability between different outcome measures of specific binding (such as V_S_ and BP_ND_), nor does it affect the evaluation of convergent validity.

The results from this study suggest that caution is warranted for applying and interpreting BP_ND_ obtained using 2TCM or BP_ND_ from kinetic models using cerebellum or SVCA4 as reference. V_T_ and V_S_ should likely be preferred over BP_ND_ from 2TCM, since they exhibited higher reliability and precision. However, the negligible correlations of V_T_ and V_S_ to SUVs are concerning and not fully understood. One explanation might be that brain SUV values are sensitive to changes in peripheral binding of TSPO (Imaizumi et al. 2007), while AIF-based outcomes are not. This hypothesis warrants further investigation in future studies. To facilitate this work, we share all TAC-data and code used in this study on a public repository (osf.io/gcn4w) for other researchers to use.

## Data and code availability

The data and code for reproducing the kinetic modelling, results, table and figure in this article can be found at osf.io/gcn4w or at https://github.com/pontusps/PK11195_TestRetest.

## Acknowledgement

We thank Jouni Tuisku from the Turku PET Center for his help with methodological considerations during the course of the study. We would like to thank the staff at the Karolinska Institutet PET centre for their assistance.

## Author contributions

PPS conceived of the study and designed the study. PPS and ZC carried out the image analysis. PPS and GJM performed the kinetic modelling. PPS carried out the statistical analyses. PPS and SC drafted the article. SC and LF supervised the study. All authors interpreted the results, critically revised the article and approved of the final version for publication.

